# Long non-coding RNA regulation of spermatogenesis and endosomal processes via the spectrin cytoskeleton in *Drosophila*

**DOI:** 10.1101/2020.07.24.220640

**Authors:** Mark J Bouska, Hua Bai

## Abstract

The spectrin cytoskeleton has been shown to be critical in diverse processes such as axon development and degeneration, myoblast fusion, and spermatogenesis. Spectrin can be modulated in a tissue specific manner through junctional protein complexes, however, it has not been shown that lncRNAs interact with and regulate spectrin. Here we provide evidence of a lncRNA that interacts with α and β Spectrin to regulate spermatogenesis and endosomal related activity in fat bodies of *Drosophila*. Protein-RNA and Protein-Protein biochemical analysis indicated the interaction between α and β Spectrin is modulated by the lncRNA CR45362. Immunocytochemistry revealed CR45362 is highly expressed in the basal testis while α and β Spectrin are clearly disrupted in this same region of CR45362 mutants. We genetically demonstrate α-Spectrin and CR45362 deficiencies cause spermatid nuclear bundling defects with congruous changes of spectrin distribution and reduced Lysotracker staining in the fat body. Our data suggests lncRNA regulation of spectrin could provide cells with a repertoire of modulatory molecules to manipulate cell-type specific cytoskeletal and endosomal requirements.

## Introduction

Long non-coding RNAs (lncRNAs) comprise roughly half of the human genome^1,2,3^ and constitute a pool of largely unexplored modulatory molecules that can regulate cytoskeletal proteins, endosomes, and autophagic pathways in disease^4,5.6^. LncRNAs also can act in a tissue specific manner^7^, in particular, they are developmentally significant as a large number of lncRNAs demonstrate high expression in the testis in wide ranging species from *Drosophila*, to mice, to humans^8,9,10,11^. The testis specificity of lncRNAs has been posited as a possible mechanism by which increased organism complexity occurs through gain of function in other tissues^7^. This specificity also offers a unique opportunity to identify targets for male non-hormonal contraceptives and identify underlying causes of idiopathic male infertility, but few lncRNAs have been mechanistically pursued in this context.

During mammalian sperm maturation, sertoli cells surround individual sperm and undergo intricate endocytic exchanges of ectoplasmic specialization and tubulobulbar complexes (actin-spectrin-clathrin structures) to migrate the maturing spermatocyte to the lumen of the seminiferous tubules and hold the spermatid until mature, all while maintaining the blood-testis barrier.^12^ In *Drosophila* the male germ cells are totally encompasses and maintained by 2 Cyst cells, which are similar to human Sertoli cells^13^. The cyst cells provide molecular cues and provide protection until sperm maturate and are released into the seminal vesicle. A key step in the sperm maturation process occurs following meiosis in which 64 spermatid nuclei are bundled tightly together within the head cyst cell; failure of this bundling process results in sterility^14,15^. Spermatogenesis requires dynamic remodeling of the actin-spectrin cytoskeleton in maturating sperm, cyst cells, and epithelial cells as the 64 sperm nuclei are brought together, condense into tight thin nuclei, elongate their tails, and remain held in a bundle within the cyst cells, which in turn have to migrate the length of the testis while interacting with the epithelial cells that make up the testis wall^16,17,18^. Although endosomal regulation of junctional complexes^19,20^ and actin dynamics^21,22^ play major roles in *Drosophila* spermatogenesis, how the complex cytoskeletal remodeling processes are tightly controlled in each interacting cell type during *Drosophila* spermatogenesis and bundling is still not completely understood. We demonstrate that the mutation in the lncRNA CR45362 gene disrupts the spermatid nuclear bundling process in Drosophila melanogaster.

Herein we identify an uncharacterized lncRNA that interacts with α and β Spectrin. The spectrin cytoskeleton is modulated in axon development and degeneration^23,24^, myoblast fusion^25^, and spermatogenesis^12^ through junctional protein complexes^26^, however, our work suggests lncRNAs can also interact with and regulate spectrin during these processes. We provide biochemical and genetic evidence that the targeted deletion mutation of the lncRNA CR45362 gene results in male sterility and alters endosomal related processes in the fat body of *Drosophila melanogaster* through α/β Spectrin disruption. This is consistent with work detailing spectrin mutants that display fat body and testis phenotypes^27,28^. Our results indicate a novel mechanism by which the spectrin cytoskeleton can be contextually regulated; though lncRNAs.

## Results

### The lncRNA CR45362 has positive effect on endo/lysosomal activity and fertility

Our lab previously performed a genome-wide association study (GWAS) to screen for genes with undescribed functions in autophagy/lysosome activity in the adult *Drosophila* fat body (unpublished). We utilized the Drosophila Genetic Reference Panel (DGRP)^29^, which consists of over 200 *Drosophila* lines that have been fully sequenced for single nucleotide polymorphisms (SNPs), and found significant autophagy/lysosome related SNPs that correlated to 152 protein-coding genes and 9 lncRNAs (Table S1). We validated our protein data using RNAi and were able to reveal proteins that had not been previously linked to autophagy/lysosome activity (unpublished). However, the lncRNA candidates could not as easily be investigated in this manner. To examine the role of these lncRNAs in autophagy regulation, we first generated CRISPR deletion mutants (of which the lncRNA CR45362, represented by CR45362KO or simply KO, will be the focus of this article) and sequenced the corresponding DNA regions to confirm successful deletions had occurred (Fig. 1A,B,S1). Additionally, we corroborated the predicted RNA length and isoform number for the CR45362 RNA in wild type flies (Fig. 1C).

**Figure 1.**
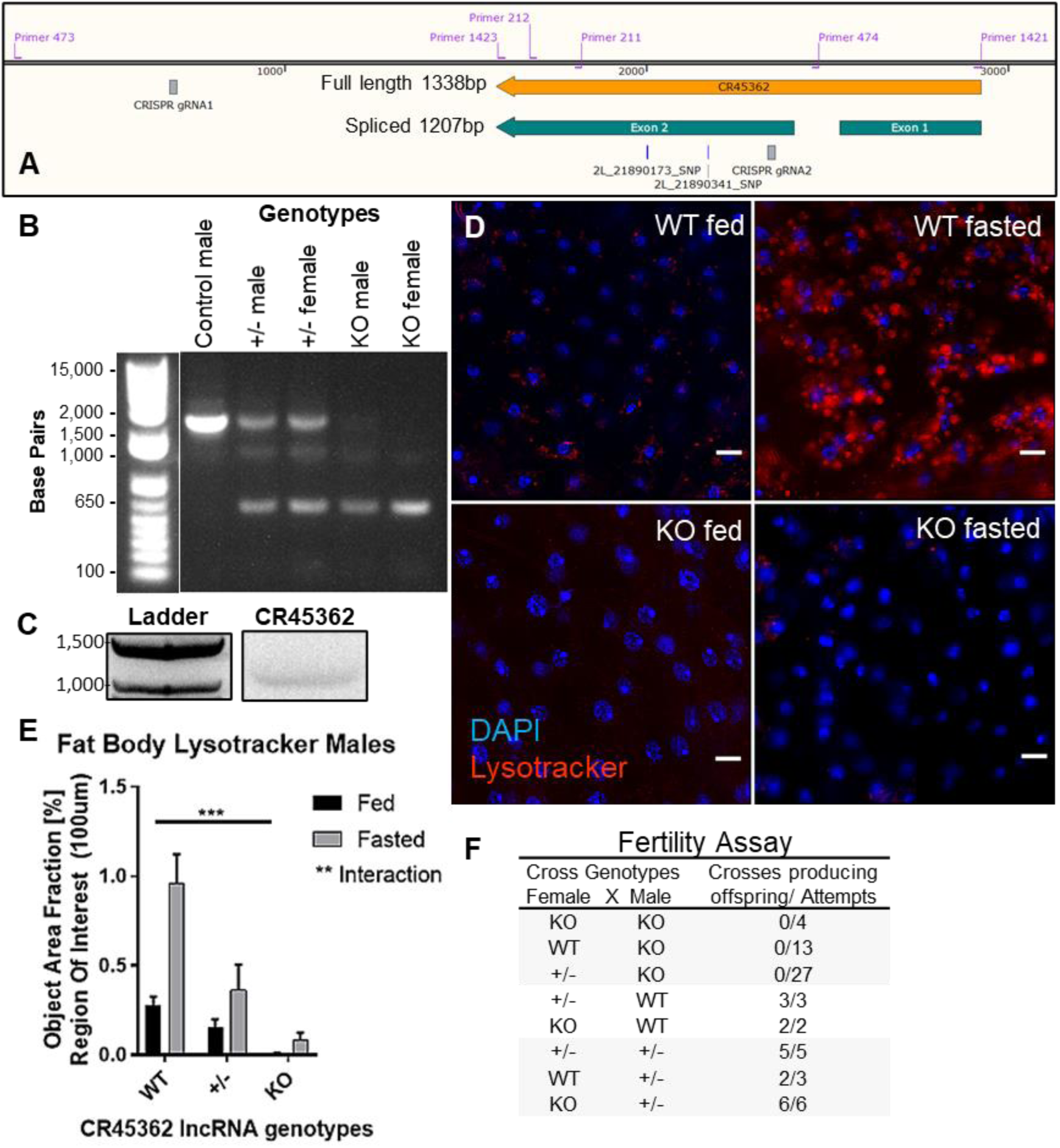
The lncRNA CR45362 has positive effect on endo/lysosomal activity and fertility. A) Diagram of CR45362, SNP’s identified in our lysosomal GWAS, primers, and CRISPR gRNAs. (Image: SnapGene 4.3.4) (B) DNA gel using primers flanking deletion target segment (p473 & 474). WT has band at 2182 demonstrating the primer targeted segment has no deletion, heterozygous CR45362 KO (+/-) has the 2182bp band but also the expected 508bp band for the deletion on the homologous chromosome, and KO has only the 508bp band indicating successful deletion in the CR45362 gene. The 508bp band was excised for sequencing and alignment (Fig. S1). (C) DNA gel of cDNA converted from CR45362 RNA using primers 1421 and 1423 (as diagrammed in 1A) that precisely correspond to the predicted ends of the lncRNA. The predicted size of CR45362 if spliced as annotated is 1207bp while unspliced is 1338bp. The gel revealed only a single band near the 1200bp mark indicating that there is a single isoform in our WT line and it is consistent with predicted splicing. Full length gel can be found in Fig. S1. (D,E) Lysotracker staining of WT and CR45362KO fed and fasted fat bodies. Knockout of the lncRNA CR45362 results in lowered Lysotracker staining under both conditions. One replicate consists of the average of three circular areas of 100μm per fly fat body measured by Cell Sens software for percentage of measured area containing fluorescence. From left to right n=7,12,6,7,6,8. Fed/fasting interaction calculated using Two-way Anova column factor (** p = 0.0034), WT vs KO fed two tailed unpaired t-test (*** p = 0.001) mean+/-s.e.m. (F) Fertility was tested by crossing individual CR45362KO male flies with WT, KO, or heterozygotes for the KO virgin females. These crosses were unable to produce offspring. However, KO and heterozygous females produced adult progeny and were fully fertile, indicating a male homozygous KO sterility defect. From top to bottom n=4,13,27,3,2,5,3,6. *All scale bars represent 10 μm. *For all figures: WT = (w1118), +/-= Heterozygous, KO = CR45362 mutant.

In order to determine if these lncRNAs do indeed have an effect on endo/lysosome activity, we next tested our lncRNA mutant fly lines utilizing Lysotracker staining in fat bodies under fed and fasting conditions to identify changes in basal (fed) and induced (fasted) endo/lysosome activity^30^. The lncRNA CR45362 had a significant (p=.0001) positive effect on endo/lysosomal activity in wild type (WT) flies compared to knockouts, while induced activity was also reduced (Fig. 1D,E).

However, during our investigation we realized that the CR45362 deletion line was difficult to maintain and had fertility issues. We therefore performed a fertility assay using simple genetic crosses (Fig. 1E) and discovered that male flies homozygous for the CR45362 deletion were unable to produce adult offspring.

### The lncRNA CR45362 localizes to endosomes in cell cytoplasm in the basal region of testis

LncRNAs function in the nucleus, cytoplasm, or even extracellularly, thus, establishing the cellular localization of a lncRNA of interest is a critical step when attempting to determine its function^31,32^. Due to the strong sterility phenotype seen in CR45362 mutants, and the lncRNA CR45362 having its highest expression by tissue type in the testis (Flybase and RT-qPCR), we performed fluorescence in situ hybridization (FISH) in the *Drosophila* testis. CR45362 had high localization to the basal region of the testis in the cytoplasm of a cell type (putative late cyst cell) that surrounds bundles of elongated spermatid nuclei (Fig. 2A,B), and it was not seen in this region of the KO (Fig. 2C).

**Figure 2:**
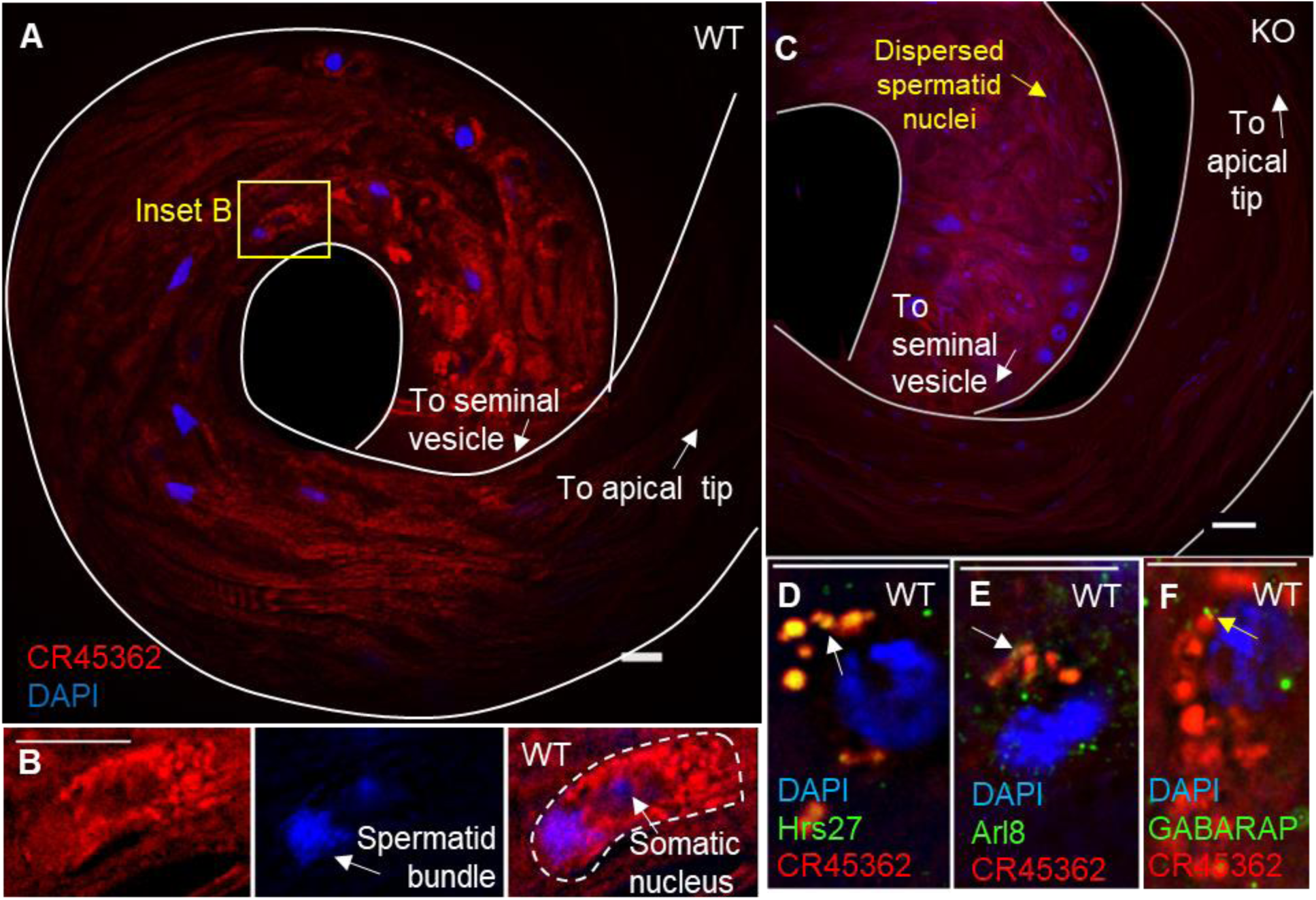
The lncRNA CR45362 localizes to endosomes and cell cytoplasm in the terminal epithelial region of testis. (A) FISH probes for the lncRNA CR45362 localize to the basal testis of WT flies. (B) Enlarged inset from Figure 2A reveals probes localize to the cytoplasm of somatic cells and a bundle of elongated spermatid nuclei can be seen infiltrating these same somatic cells. (C) Basal testis in KO for same region as Figure 2A shows no probes, and indicates successful knockout. (D) CR45362 probes co-localized (white arrows) with antibodies against the early endosome marker Hrs but were less often localized with lysosome marker arl8 (E). Probes did not colocalize with autophagy marker GABARAP but were occasionally near this marker (yellow arrows) (F). *All scale bars represent 10 μm.

Given that CR45362 was identified in a lysosome GWAS and mutation of the gene affects lysosome activity in the fat body, we combined FISH with Immunohistochemistry to determine if this function may be conserved in the testis. Interestingly, CR45362 probes co-localized with antibodies against the early endosome marker Hrs27 (Fig. 2D) but infrequently with lysosome marker Arl8^33^ (Fig. 2E). Probes did not colocalize with autophagy marker GABARAP (Fig. 2F), a *Drosophila* atg8 homolog^34^, but were occasionally seen near this marker.

### Disruption of CR45362 results in a loss of spermatid nuclear bundling

Although FISH indicated that CR45362 is expressed in late stages of spermatogenesis, it is still critical to eliminate known early stage defects as they often phenotypically replicate defects that manifest during later stages^15,35,36^. We used immunostaining or phase contrast imaging to examine the hub cells (Fig 3A), round stage spermatids with nebenkern (Fig. 3B), and canoe stage spermatids (Fig. 3C). All appeared normal. We backcrossed the CR45362 mutation into dj-GFP lines and stained the nuclei with DAPI, which revealed that the spermatid tails were developed and spermatid nuclei were elongating (Fig 3D). However, the CR45362 mutation was disrupting the bundling of the 64 elongated spermatid nuclei (Fig. 3D,E) which were unable to enter the seminal vesicle (Fig. 3F). Permeability assays revealed that the defect was not due to the cyst cells detaching from each other, as spermatids remained encapsulated by the cyst cells (Fig. 3G,H). An important step in sperm development that occurs shorty after nuclear bundling in *Drosophila* is individualization. An individualization complex of actin “cones” move the length of the spermatids and remove cytoplasmic contents. In CR45362 mutants the actin cones are dispersed because the spermatid nuclei were never able to bundle together before the actin cones formed (Fig. 3E). Serendipitously, dispersed actin cones and elongated nuclei serve as ideal phenotypes for genetic screening as phalloidin (actin) and DAPI (nuclei) staining can be performed in less than one hour while being readily identifiable. The combined imaging demonstrated that the sterility defect was occurring after elongation of the spermatid nuclei and tails, but before spermatid individualization.

**Figure 3:**
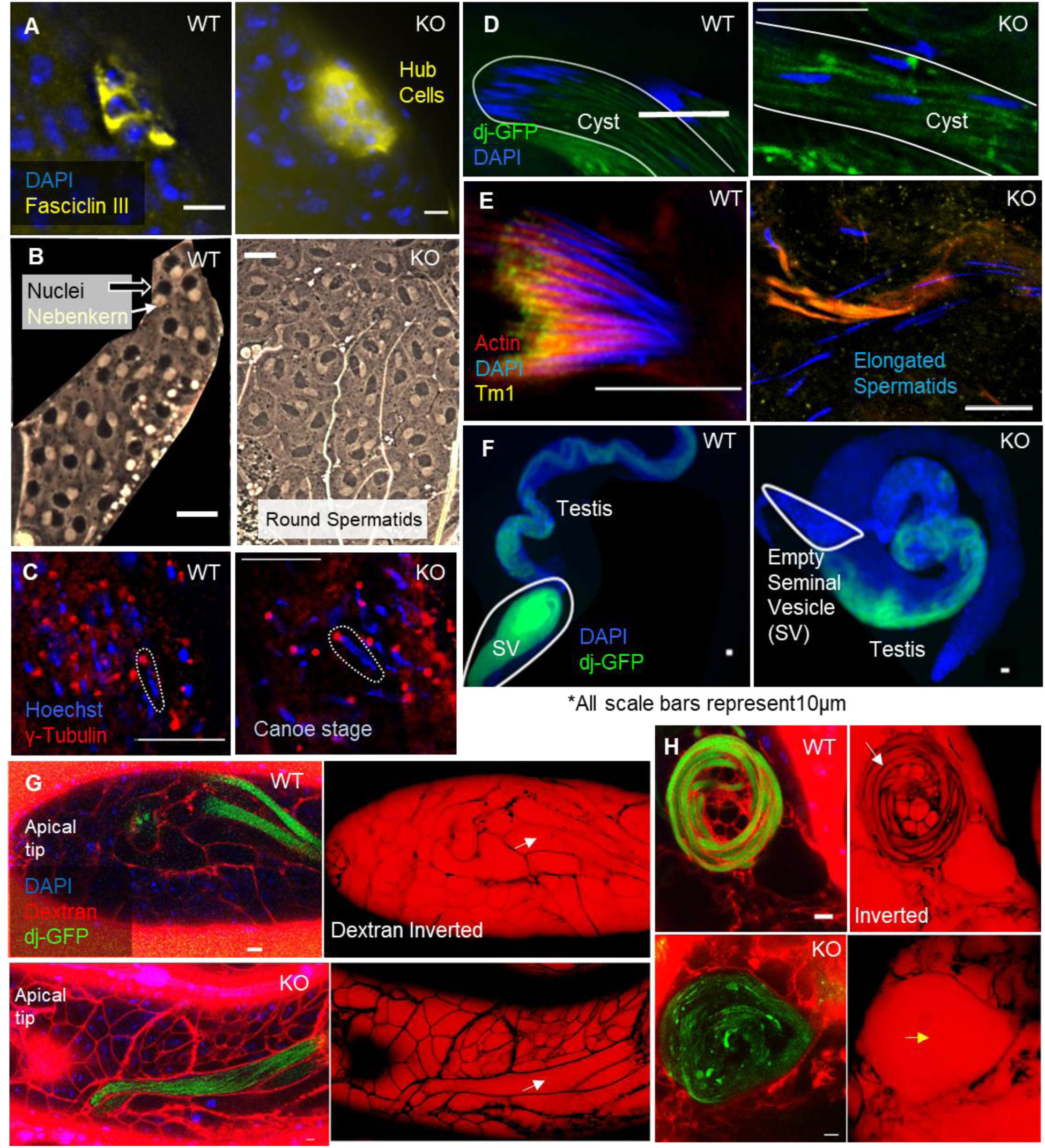
Spermatid nuclear bundling process impaired in KO lines. (A) Hub cells indicated by Fasciclin III have proper radial arrangement^55^, position at the apical tip^56^, and number^57^. (B) Phase-contrast imaging of wild-type and KO spermatids show a 1:1 ratio of nuclei (black arrow) to nebenkern (white arrow), indicating no defects in round spermatids due to failed cytokinesis, errors in chromosome segregation, nebenkern fusion, etc.^58^. (C) γ-tubulin marks the centriolar adjunct^59^. Centriolar adjuncts are associated with spermatid nuclear scattering phenotypes in which γ-tubulin is not localized near the nuclei. Our immunostaining indicated proper attachment in KO^14,60^. Dotted lines highlight a single nuclei and properly attached basal body. (D) Spermatid flagella form, marked by dj-GFP^61^, and nuclei are elongated, however nuclei are dispersed throughout the cyst in KO. (E) Actin cones form (indicated via Phalloidin staining) and are decorated along their length by Tm1 in WT and KO spermatids. Indicating the defect occurs independent of individualization, as spermatids are dispersed before actin cone formation^62^. (F) Spermatids (indicated by dj-GFP) fail to enter the seminal vesicle in KO males. (WT testis were manually uncoiled for imaging) (G) Permeability assay using 10kD Dextran dye indicates that the cyst cell permeability barrier surrounding elongating spermatids is intact (white arrows) as don juan GFP reveals spermatid tails that appear surrounded by un-ruptured tail cyst cell^63^. (H) Impermeable coiling stage cysts (yellow arrow) in KO indicate proper cyst cell enclosure. Cyst, likely in stages of spermatid release (white arrow) has loss of permeability barrier in WT. *All scale bars represent 10 μm.

### The lncRNA CR45362 biochemically interacts with cytoskeletal and endosomal proteins

The cytoplasmic localization of CR45362 indicated that this lncRNA was not regulating transcriptional processes in the nucleus, while the co-localization with endosomal markers suggested the action of CR45362 could be through interaction with proteins, therefore, to determine the protein binding partners for CR45362 we performed Chromatin Isolation by RNA Precipitation – Mass Spectrometry (ChIRP-MS). ChIRP-MS utilizes reversible protein-lncRNA crosslinking followed by magnetic bead pulldown of biotinylated probes complementary to the lncRNA of interest (Fig. 4A). Reversing the crosslinking and then performing Mass-spectrometry identifies candidate proteins that bind to the lncRNA. Gene Ontology (GO) Molecular Function analysis for the 50 most highly enriched proteins had 2 groups stand out: cytoskeletal and GTP binding groups (Fig. 4B). This was significant because the enriched cytoskeletal proteins contained components of actin-spectrin junctional complexes^37^, while the GTPase category is a group of proteins considered the “master regulators of intracellular membrane traffic” ^38,39^. Most importantly, the results of the ChIRP-MS (Table S4, Fig. 4E) supported our FISH experiment outcomes in that the proteins identified were cytoplasmic and endosome related proteins.

**Figure 4.**
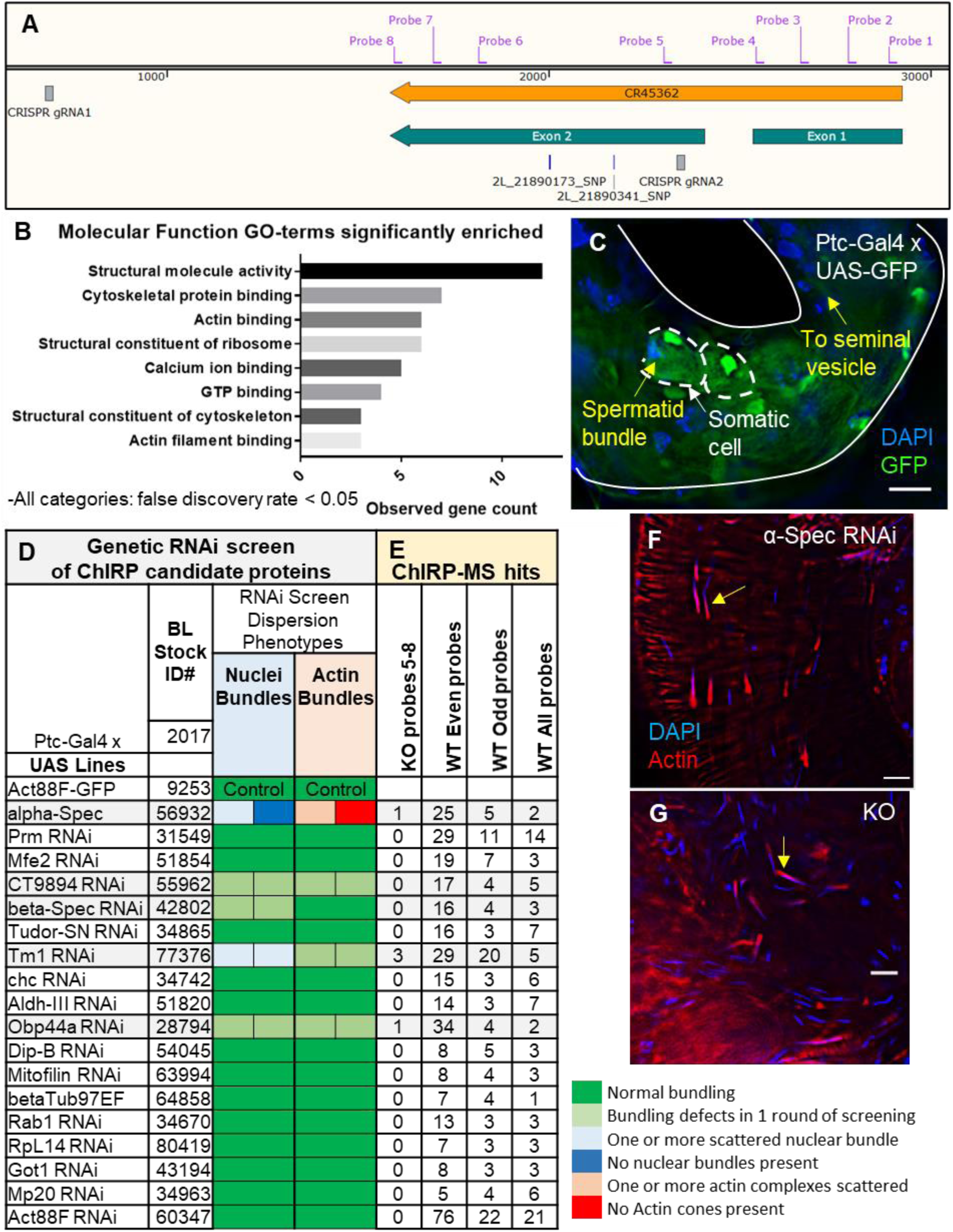
The lncRNA CR45362 biochemically interacts with cytoskeletal and endosomal related proteins. (A) Map of ChIRP probe locations relative to CR45362 gene. Probes 5-8 map to the deleted segment of the gene in the KO. (Image: SnapGene 4.3.4) (B) Molecular Function Gene Ontology for top ChIRP-MS proteins indicates cytoskeletal proteins bind CR45362. (C) Driver test indicates Ptc-Gal4 x UAS-GFP expresses GFP in the same region of the basal testis as is indicated by CR45362 FISH (see also Fig. 2A). (D) RNAi screen of top ChIRP-MS candidate proteins. Five pairs of testis from 3-5 day old males from each UAS-Gal4 cross were dissected of which we scored 3. If defects were seen, we repeated this screening for a second round of crosses. If both testis of a pair had the same phenotype, they were scored according to the following criteria: normal nuclear bundling, one or more scattered nuclear bundles, no nuclear bundles present, one or more individualization complexes (actin) scattered, no actin cones present, or testis deformed. (E) Results from ChIRP-MS. Columns list the number of hits (spectra) for each protein. Numbers are internally relative to each individual sample, and are used for ranking proteins, cut-off selection, and identifying background proteins. Probes 5-8 (Fig. 4A, S3) cover the CRISPR excised region, when mixed with KO samples these probes serve to identify background proteins pulled down by beads, general biotin probes, and as a negative control for the pulldown of CR45362 (Fig. S4). Each individual probe could have probe-specific background, therefore established protocols54 recommend the optimal control for a ChIRP assay is to pool probes into “Even or Odd” groups in WT samples, allowing further identification of background proteins and can be compared with the WT sample mixed with “All” probes. (F) The Ptc-Gal4 x α-Spec RNAi line has dispersed actin cones (indicated by phalloidin staining) and spermatid nuclei (yellow arrows), which phenocopied the nuclear dispersion seen in CR45362KO (G). *All scale bars represent 10 μm.

### Alpha Spectrin RNAi leads to nuclear bundling defect

Using the spermatid nuclear bundling and actin cone dispersion phenotypes found in our investigation above, we performed a UAS/Gal4 RNAi screen of 20 top proteins from the ChIRP-MS utilizing the UAS-Gal4 system (Fig. 4D). Although we examined 8 different testis specific Gal4 driver fly lines^40,41^ (Table S3), we performed the screen using a driver line with GAL4 under the control of ptc (Bloomington #2017). When crossed with UAS-GFP, this line expressed GFP in the basal testis in a pattern similar to the FISH results for CR45362 (Fig. 4C, 2A). The genetic screen revealed that α-Spec RNAi (Bloomington Stock #56932) crossed to the ptc-GAL4 line had the most similar nuclear bundling defect to that of the CR45362 mutant. Both α-Spec RNAi and CR45362 mutants had elongated spermatids decorated with actin cones but these were dispersed throughout the cyst (Fig. 4F,G). The knockdown efficiency of the α-Spec RNAi was likely limited as some nuclear bundling did in fact occur and there were weakly assembled individualization complexes present. Four other proteins from the top ChIRP-MS candidate list β-Spec, Tm1, CT9894, and Obp44a had weaker nuclear bundling defects under this GAL4 line, but only Tm1 RNAi passed both rounds of screening and the effects were less than that of α-Spec RNAi. It is important to note that α-Spec interacts with β-Spec and tropomyosin (via actin) at spectrin junctional complexes^37^. Therefore, these results suggested a genetic interaction with CR45362, and supported further interrogation of α-Spec.

### Alpha & Beta Spectrin disrupted in basal testis of CR45362 mutant

We next asked, what effect does the mutation of CR45362 have on spectrin in the testis? To answer this we performed immunostaining against α-Spec in the testis of WT and CR45362 mutant lines. We saw that fusomes formed and elongation complexes were still present in the apical and medial testis of the mutant respectively (Fig. S5). However, in the basal testis α-Spec was dispersed throughout the somatic cells of the KO (Fig. 5A-D). During our previous immunostainings we had not seen this type of dispersion in these cells before and it indicated that there was a clear disruption of normal α-Spectrin in the same region of the testis as CR45362 was seen during in situ of WT testis.

**Figure 5.**
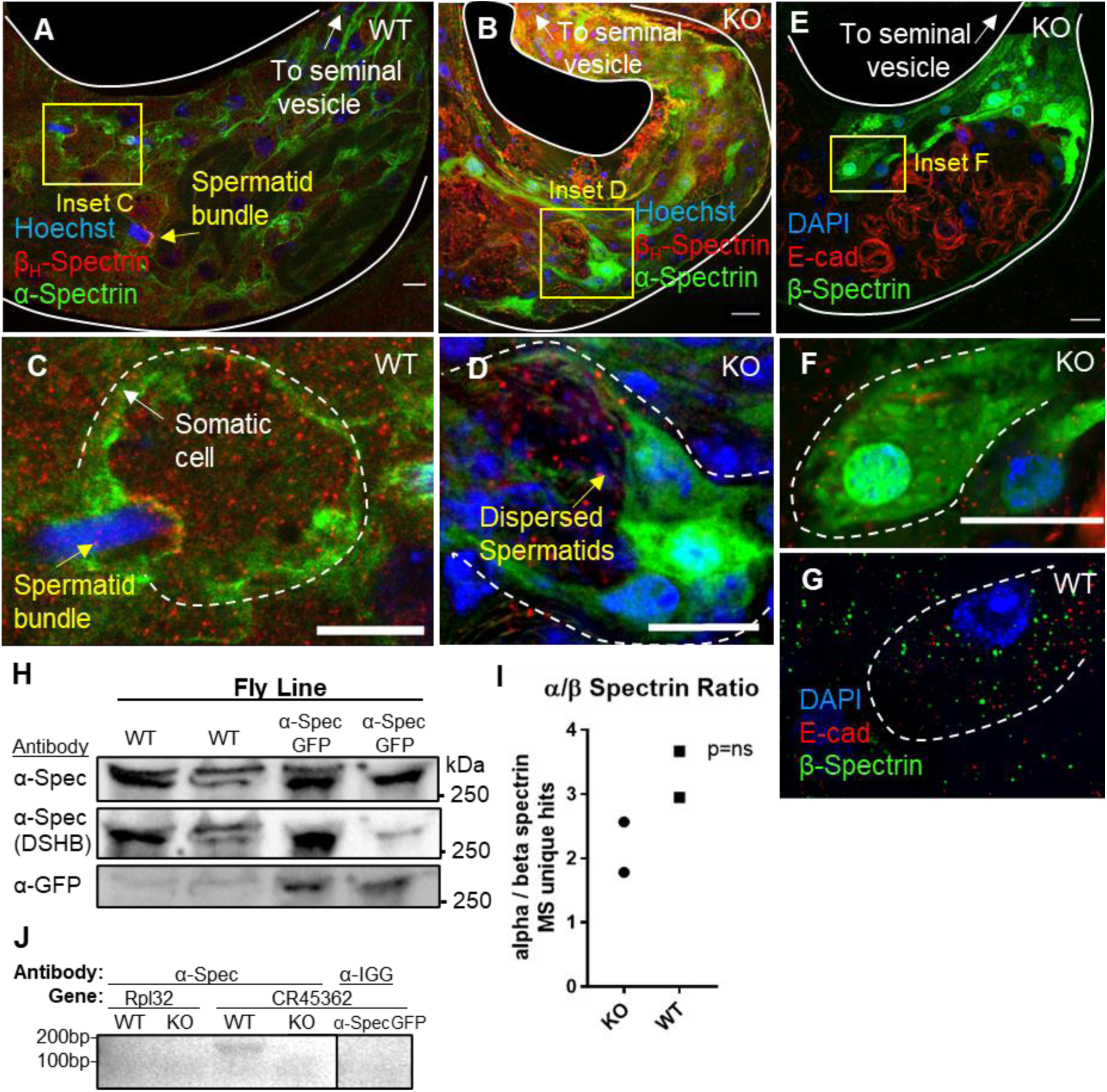
Alpha & Beta Spectrin disrupted in basal testis in CR45362 mutant. (A,C) Under immunostaining α-spectrin localizes to the exterior of the somatic cells in the basal testis, while Beta Heavy Spectrin accumulates in the somatic cell at the location where the spermatid nuclear bundles are anchored. (B,D) Alpha Spectrin is dispersed throughout somatic cells in KO with enhanced nuclear membrane localization in the cyst cell. (E) Beta Spectrin in the basal testis of the KO has a dispersion pattern similar to α-Spectrin in KO testis. (F) Inset of enlarged somatic cells showing dispersion of β-Spectrin in KO vs WT (G). (H) Western blots for α-Spectrin antibodies. First 2 columns are WT replicates, columns 3 and 4 are from an α-Spectrin fused GFP line (Table S3). Two α-spectrin antibodies were tested (rows) and have bands at the same MW as the α-GFP antibody (third row). The image is of the same location after stripping the membrane between each separate antibody incubation. ∼Size estimates based on 250kDa standards band and 278 kDa MW of α-Spectrin (full-length blots, Fig. S6) (I) Co-immunoprecipitation for α-Spectrin indicates the ratio of α-Spectrin to β-Spectrin increases in WT testis. n=2 Unpaired t-test (p = 0.1678, ns = not significant) (J) Agarose gel from α-Spec RIP-PCR. Antibodies against α-Spec pulled down CR45362 RNA (CR45362 primers 211-212 product size = 142bp) but not Rpl32 in WT testis. IgG control antibody did not pull down CR45362 in α-Spec-GFP fly lysate. Full length gels can be found in Fig. S7. *All scale bars represent 10 μm.

The evidence was implicating the spectrin junctional complexes as being somehow regulated by the CR45362 lncRNA. Consequently we wanted to determine if the composition of the α-Spec junctional complex was changing in CR45362 mutants. We first validated two α-Spec antibodies (a generous gift from the Dubreuil lab vs. a Developmental Studies Hybridoma Bank - DSHB - antibody) by performing westerns with an α-Spec-GFP line compared to WT (Fig. 5H, S6). We then dissected KO and WT testis to carry out Co-Immunoprecipitation Mass Spectrometry using the DSHB α-Spec antibody. Of the proteins pulled down with α-Spec, two proteins stood out; Hsp27 which was in the KO but not in WT, and β-Spec (Table S4). Importantly, β-Spec was the most highly enriched of all proteins not pulled down by the control beads (lysate with no antibody), but also the ratio of α-Spec to β-Spec showed an increase in WT vs mutant testis (Fig. 5I). Although not strictly quantitative, or reaching levels of significant due to low replicate numbers (n=2, from 1,240 dissected testis), there was a change from 1.8-2.6 α-Spec to β-Spec ratios in CR45362 mutant samples compared to 2.9-3.6 ratios in α-Spec to β-Spec in WT. This is of note because it has been shown that the ratio of α-Spec to β-Spec is critical to cell function^28^. Hsp27 was not significant in our ChIRP so we performed immunostaining against β-Spec. The KO testis exhibited a dispersion of β-Spec (Fig. 5E-G) in the same region in which we had previously seen α-Spec dispersion and CR45362 localization. These results provided further biochemical evidence that the interaction between α and β Spec is being regulated by CR45362, and are consistent with nuclear bundling defects seen in our genetic α and β Spec RNAi screen.

To confirm that α-Spec was interacting with CR45362 we extracted RNA from the WT α-Spec Co-IP pulled-down material. We were able to detect CR45362 RNA in RT-qPCR but only very weakly (Fig. S7). We therefore dissected more testis and performed a RNA immunoprecipitation (RIP), where RNA-protein crosslinking is performed followed by α-Spec antibodies being used to pull down α-Spec protein complexes, de-crosslink, isolate RNA, and then perform PCR. RT-qPCR results indicated that CR45362 was present in WT but not KO samples and there was no housekeeping RNA (Rpl32) present in either sample, indicating spectrin was binding CR45362. RT-qPCR detection levels were low so we ran an agarose gel and were able to see a band for CR45362 in WT (Fig. 5J). We were unable to detect CR45362 using α-GFP or IgG in samples from an α-Spec-GFP fused line (Fig. S7).

### Alpha Spectrin knockdown and CR45362KO alter fat body endosomes and trafficking

The CR45362 lncRNA was identified through a fat body lysosome GWAS, thus we asked if regulation of spectrin could also explain the fat body Lysotracker phenotype seen in CR45362 mutants. We crossed a fat body Gal4 driver line (Bloomington# 33832) to the UAS α-Spec RNAi line used in our aforementioned nuclear bundling screen, and then stained the adult fat body with Lysotracker. For this Lysotracker assay we additionally backcrossed the KO into an alternative YwR background to reduce possible confounding genetic elements and further rigorously confirm our initial Lysotracker results. As predicted, both α-Spec RNAi and CR45362 mutant fat bodies had reduced endo/lysosome activity (Fig. 6A).

**Figure 6.**
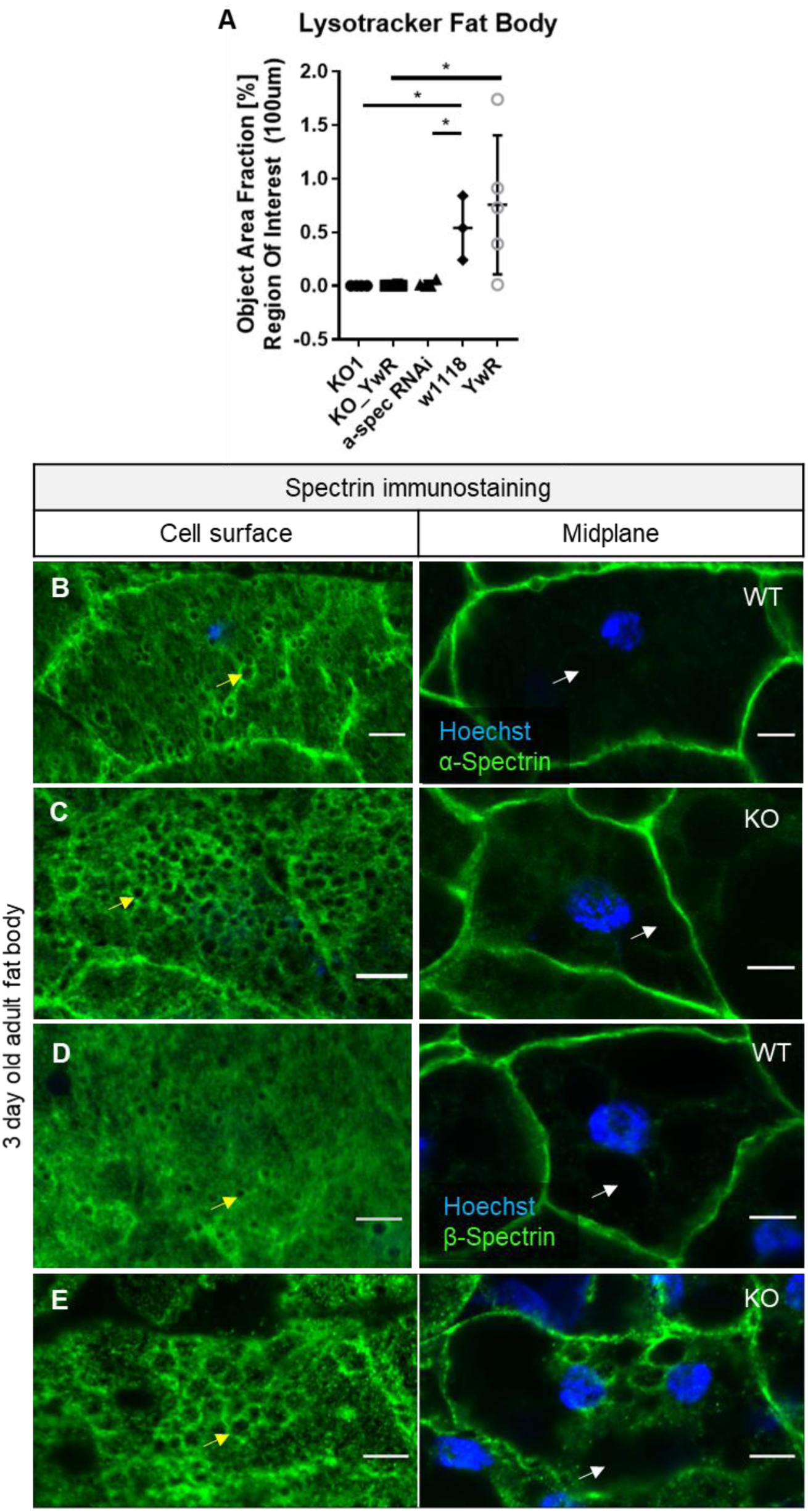
Alpha Spectrin knockdown and CR45362KO alter fat body endosomes and trafficking. (A) Lysotracker assay comparing 5 day old fat bodies in 2 KO lines (KO1, and KO backcrossed into YwR control line = KO_YwR), fat body specific α-spectrin RNAi, and 2 WT lines (w1118 & YwR). Two-tailed unpaired t-test (*** p < 0.001, ** p < 0.01, * p < 0.05, ns = not significant) mean+/-s.e.m. (B-E) Fat bodies immuno-stained for α-Spec (B,C) and β-Spec (D,E) in 3 day old adult fat bodies of WT and CR45362KO. Cell surface images are in the left column and the mid-plane of the same cell is in the right column. Yellow arrows indicate surface droplets, white arrows indicate medial putative lipid droplets. *All scale bars represent 10 μm.

Fat bodies have 2 populations of lipid droplets with distinct regulatory networks; surface lipid trafficking dependent or centrally localized large internally generated lipid droplets, of which spectrin is involved in regulating the surface lipid trafficking-generated droplet pools^28,42^. To determine if spectrin is altered in CR45362 fat bodies we immuno-stained for α and β Spectin (Fig 6B-E) in adults. We saw large dark cavities (consistent with fat body internal lipid droplets) at the midplane that were not surrounded by spectrin. Near the membrane however, spectrin clearly encircled the dark droplets. This was consistent with spectrin’s role in lipid trafficking at the cell surface^28,42^. Interestingly, the surface of the WT fat bodies had a different spectrin pattern from KO fat bodies. Both α and β spectrin encircled the droplets at the surface but the droplets increased in size (not quantity) in the KO flies and had an altered distribution, further indicating spectrin-mediated trafficking is indeed altered in the fat bodies of CR45362KO flies.

## Discussion

We have demonstrated that the lncRNA CR45362 has a novel interaction with spectrin and that mutating the CR45362 gene results in α/β Spectrin disruption in cells of the basal testis in *Drosophila melanogaster*. Additionally, knockdown of α-Spec or mutation of CR45362 both result in loss of spermatid nuclear bundling, affect endo/lysosome activity, as well as alter surface droplets in the fat body. While our protein-protein data indicate that CR45362 is likely important for α/β Spectrin interaction.

BH-Spectrin has been shown to be necessary for Rab5 early endosome formation and functional H+ V-ATPase recruitment in the apical brush border of midgut epithelia^43^, therefore one might predict that α and β-Spec interaction would also have endosomal effects. Indeed, “Spectrin/ankyrin-G domains exclude clathrin and clathrin-dependent cargo, and inhibit both receptor-mediated and bulk endocytosis.” ^44^ It is possible that the competition of β vs βH-Spectrin interactivity with α-Spec is affected by CR45362 interaction with β-Spec, as the B isoforms tend to be mutually independent in their functions such as polarization and membrane deformational effects^45^. CR45362 may alter this isoform competition by regulating β-Spec to α-Spec binding, thus reducing early endosomal activity. Therefore, the restriction of early endosomes could also result in restriction of lysosomes^46^. Another intriguing possibility could be that Spectrin’s E2/E3 enzymatic activity is regulated by lncRNA to alter β-Spec isoform interaction, as it has been shown lncRNAs can affect ubiquitination activity^47,48^.

Meanwhile, actin and specialized junctions are known to hold the sperm nuclei to the cyst cell^19,21^, but how the nuclei are drawn together into a tight bundle is still unknown. Similar to forces involved in the process of myoblast fusion where “Myoblast invasion triggers mechanosensitive accumulation of βH-Spectrin” ^25^ we see accumulation of βH-Spectrin at the site of spermatid penetration into the cyst (suggesting the actin cap may be similar to the brush border of the midgut). And just as “a pool of free α /βH-Spectrin heterotetramers” ^25^ is generated once the forces are removed in myoblast fusion, we see a dispersion of spectrin throughout the somatic cells in CR45362KO testis. This is possibly due to the lack of proper force generation or membrane deformability that would allow the cyst membrane to sense and fold around a bundle of spermatid nuclei.

The high expression levels of lncRNAs in the testes^10,49,50^ has been posited as a possible mechanism by which increased organism complexity occurs through gain of function in other tissues^7^. This aligns well with the correlation of the increasing number of lncRNAs with increasing organism complexity^7^. Adding to this reasoning, we demonstrate that lncRNAs are able to support structural tasks in cells and that lncRNAs may play a significant role in the modulation of cytoskeletal structures which are also known to contribute to the increasing tissue complexity of higher organisms^51^. LncRNAs have been identified as modulating actin and intermediate filament cytoskeletal components in cancer cell cultures^4^, and our current spectrin in vivo experimental data further extends this overall lncRNA-cytoskeletal framework.

This work provides unique data indicating that the spectrin structures in the testes and fat bodies can be organized by lncRNA. This creates several new questions that provide interesting opportunities for future work. First, is spectrin regulated by lncRNAs during mammalian spermatogenesis, and what RNA motifs regulate their interaction? Does the spectrin cytoskeleton have a diversity of tissue specific regulatory lncRNAs, and how do these lncRNAs influence endocytic processes in those tissues? With humans having many testis specific lncRNAs, spectrin related lncRNAs could be viable targets for contraceptive development or infertility treatment, particularly with the advent of successful Antisense Oligonucleotides by Ionis Pharmaceuticals and other promising paths in the oligonucleotide field^52^.

## Methods

### Generation of knockout lines

Knockouts were generated using previously described protocols https://flycrispr.org/target-finder/, briefly, 2 CRISPR gRNAs (Table S2) were designed using CRISPR Optimal Target Finder^53^ for each lncRNA. pU6-2-gRNA plasmids (*Drosophila* Genomics Resource Center stock# 1363) were amplified in One Shot Top 10 competent E.coli cells (Invitrogen# C404010) on SOC Medium (Corning# 46003CR), run on agarose gel, and isolated using Promega Wizard Plus Miniprep DNA Purification (Promega# A1460). The pU6-2-gRNA plasmid was cut with BBSI-HF restriction enzyme (Fisher Scientific Catalog# NC1222468) in 1X CutSmart Buffer and de-phosphorylated with Calf Intestinal Alkaline Phosphatase (Fisher# 50811712). gRNA oligos were annealed using T4 Polynucleotide Kinase (Fisher# 50811602) in T4 ligation buffer (New England Biolabs# B0202S) and then inserted into pU6-2-gRNA plasmids using T4 DNA ligase (New England Biolabs # M0202S).

Plasmids containing the gRNAs were then amplified, run on gel, and isolated as described above and sequenced at the Iowa State University DNA Facility on the Applied Biosystems 3730xl DNA Analyzer. Sequences were aligned using SnapGene software (from Insightful Science; available at snapgene.com). gRNA plasmids were injected into eggs containing the Cas9 enzyme (Rainbow Transgenic Flies INC.# 51324) by Rainbow Transgenic Flies Inc. The CR45362 deletion was maintained by crossing in a cyo balancer (from line yw;Sp/Cyo;TM2e/TM6BTbHue from the Tatar lab).

## Lysotracker Assay

Flies were raised at 25^°^C on vials of food mix: 1700ml water, 15.8g AGAR (USB Cat#10654), 50g YEAST (Genesee scientific Cat#62-108), 104g CORNMEAL (Fisher scientific Cat#NC9349175), 220g commercial granulated sugar, 4.76g TEGOSEPT (Genesee scientific Cat#20-258) and 73.6ml 95% ethanol. To induce starvation response 3-5 day old male flies were put on 1ml of 1X PBS (Life Technologies # 10010-023) added to a Kimwipe in vial overnight for 12-16 hours. Fat bodies were dissected in 1X PBS then incubated in 1XPBS + Lysotracker (Invitrogen # L7528) + Hoechst (Immunochemistry Technologies# 639) according to manufacturer’s specifications for 3 minutes and imaged. Three circular areas of 100μm were measured for each fly fat body and then averaged to count as one data point, with multiple flies used for each genotype. Cell Sens software was used to quantify the percentage of measured area containing fluorescence. Significance was calculated on GraphPad PRISM version 7.03 using unpaired t-test and for Fig.1 fasting interaction was calculated using Two-way Anova column factor. All graphs were generated using GraphPad PRISM version 7.03.

### Polymerase Chain Reaction

RNA was extracted using Trizol (Life Technologies #15596026) and subjected to DNase treatment (Turbo DNA-free, Ambion# AM1907). cDNA was made using the qScript cDNA Synthesis Kit (Quanta Bio #95047-100) or using the First Strand cDNA Synthesis Kit (NewEngland Bio# E6560) using the random primer mix for full length CR45362 following manufacturer’s protocols. PCR was performed using manufacturer’s protocols from DreamTaq DNA Polymerase (Thermo Scientific # EP0702) and run on a 2% agarose gel, while quantitative PCR was performed using PowerUP SYBR Green Master Mix (Thermo Fisher# A25742).

### Fluorescence in Situ Hybridization

RNA from WT 3-5 day old flies was extracted and reverse transcribed to cDNA using Power SYBR Green Cells-to-Ct Kit (Invitrogen# 4402953) following manufacturers protocol. Oligos were generated following manufacturers protocol by PCR Amplification using DreamTaq DNA Polymerase with buffer (Thermo Scientific# EP0701), 10mM dNTPs (Sigma Aldrich# DNTP100-KT), 5pmol/ul custom primers #427 and #212 (Table S2), and ∼1ug of cDNA from above. Products were run on agarose gel and purified using PureLink Quick Gel Extraction Kit (Invitrogen# K2100-12). Oligos were then amplified using pGEM-T Easy Vector kit (Promega# A1360), One Shot Top 10 competent E.coli cells (Invitrogen# C404010), SOC Medium (Corning# 46003CR), purified with Promega Wizard Plus Miniprep DNA Purification (Promega# A1460), and then sequenced at the Iowa State University DNA Facility on the Applied Biosystems 3730xl DNA Analyzer to confirm proper oligo insertion. Plasmids were linearized using NcoI (New England Bio Labs# R0913S) and BspQI (New England Bio Labs# R0712S) Restriction enzymes in NEBuffer 3.1 (New England Bio Labs# B7203S) according to manufacturer’s specifications and purified using PureLink QuickPCR Purification Kit (Catalog# K3100-01). Fluorescent probes were generated following the Invitrogen FISH tag RNA Red Kit (Invitrogen# F32954) manufacturer’s protocol. Materials used but not included in the kit are as follows: Xylene (Fisher# X5-500), Proteinase K (Invitrogen# 25530-049), Formamide (Fisher#BP227-100), Heparin Sulfate powder (Alfa Aesar# A16198), 20X SSC (Invitrogen# AM9763), Herring testes DNA (Promega#D1811), Tween-20 (Fisher# BP337-500), 5% Normal Donkey Serum. Following post-hybridization steps, samples underwent protein immunostaining (see Methods; Immunocytochemistry below).

### Immunocytochemistry

The following protocols are dependent on antibodies and samples. Testes from 3-5 day old flies were dissected in 1X PBS, Fixation in 4% Para-formaldehyde diluted in 0.1-0.3% PBST (1X PBS plus Triton X-100, Fisher# BP151-500) for 20-30 mins at room temperature. Samples were washed and optionally blocked with PBST + BSA (0.25g Bovine Serum Albumin (Sigma Aldrich# A7906-500G) per 50ml PBST) three times, 10 mins each. The samples were incubated with Primary antibody overnight at 4 °C or for 2hrs at room temperature. Samples were next washed in PBST 3 times for 10 minutes each wash, then incubated with secondary antibodies (1ul secondary + 400ul PBST) for 2 hours at room temperature. Samples were again washed in PBST 3 times for 5 minutes each. Samples were either stained with 1ul Hoechst (Immunochemistry Technologies# 639) + 5ul Alexa Fluor Phalloidin - if actin staining (Thermo Fisher scientific# A12381) + 200ul PBST for 30 mins followed by one wash in 1X PBS and mounted in ProLong Diamond Antifade Mountant (Invitrogen# P36961). Alternatively this staining was replaced with mounting in Prolong Gold anti-fade reagent with DAPI (Invitrogen# P36939). Primary antibodies are as follows; DSHB antibodies were used at concentrations of 2-5ug/ml. Arl8 (DSHB # arl8-p), Hrs27 (DSHB# Hrs 27 – 4 -s), E-Cadherin (DSHB #5D3), Fasciclin III (DSHB # 7G10), α-Spectrin (DSHB # 3A9 (323 or M10-2)-s), γ-Tubulin (1:500) (Sigma Aldrich# T5326), GABARAP (1:300) (Cell Signaling Technologies# E1J4E), α-Spectrin (1:2000) and β-Spectrin (1:1000) were generous gifts from Dr. Ronald Dubreuil at UIC, βH-Spectrin (kst) (1:1000) was a generous gift from Dr. Claire Thomas at PSU. Secondary antibodies (1:500); α-Rabbit (488) (Jackson ImmunoResearch # 711-545-152), α-Rabbit (594) (Jackson ImmunoResearch # 711-585-152), α-Rat (647) (Jackson ImmunoResearch # 712-585-150), α-Mouse (488) (Jackson ImmunoResearch # 115-545-166), α-Mouse (594) (Jackson ImmunoResearch # 115-585-166)

### Permeability Assay

Testis were dissected from 3-5 day old flies and incubated in Corning Insectagro DS2 Serum-Free/Protein-Free Medium (200ul medium + 1ul Hoecsht) for 5 minutes, transferred to 15ul of 10 kDa dextran conjugated Alexa Fluor 647 Dextran AlexaFluor 647 (Invitrogen# D22914) at 0.2 μg/μl, and imaged within 60 minutes.

### Microscopy and image analysis

Olympus BX51WI fluorescent microscope and Olympus IX83 confocal microscope images were processed with Cell Sens imaging Software and finalized using FIJI. Phase contrast images were taken on a Zeiss Axioskop phase contrast microscope capture with ProgRes MAC CapturePro 2.7.

### Chromatin Isolation by RNA Precipitation – Mass Spectrometry

ChIRP-MS was performed using published protocols^54^. AntisenseDNA Oligos were purchased from, and designed (Fig. S2), using Bioresearch Technologies ChIRP Probe Designer at: https://www.biosearchtech.com/support/tools/design-software/chirp-probe-designer. Testis from 373 - 662 flies (3-5 day old) for 3 WT replicates were performed using even or odd tiling of probes, or all probes (Fig. 4A, S3). Probes corresponding to the deleted segment were used as a control in the CR45362KO. Testis were dissected in ice cold RNase free 1X PBS with tools and surfaces cleaned by RNaseZap RNase Decontamination Solution (Ambion# 9780), fixed in 3% pfa, quenched with 2.5M glycine (Sigma Aldrich# 410225-50G), washed in 1X PBS, flash frozen, and stored at -80. All ChIRP-MS steps were followed as described in Chu and Chang 2018. RT-qPCR confirmed that the probes pulled down CR45362 but not the RpL32 control (Fig. S4). Mass spectrometry was performed by The Taplin Biological Mass Spectrometry Facility at Harvard University. STRING-db https://string-db.org/ was used for Gene Ontology analysis. Results from each sample were sorted by number of hits. Proteins in WT samples that had over 3 hits in KO samples were excluded. Proteins in WT samples that had 2-3 hits in KO samples were excluded unless they had over 5 hits in both even and odd probe WT samples. Proteins in WT samples that had only 1 hit in KO samples were excluded unless they had over 4 hits in both even and odd probe WT samples. The “All probes” trial had increased diversity of probes causing numbers of protein species to increase, which in MS, results in fewer hits detected per protein species but were still included in analysis. All graphs were generated using GraphPad PRISM version 7.03.

### Genetic RNAi screen

We used phalloidin and DAPI (described in Methods: Immunocytochemistry) to screen for spermatid nuclear dispersion and dispersed actin cones. We first screened all UAS and Gal4 lines (Table S3) without crosses to ensure that there were no background effects on nuclear bundling. Gene names were replaced with numbers to prevent bias. We next dissected 5 pairs of testis from 3-5 day old males from each UAS-Gal4 cross of which we scored 3. If defects were seen, we repeated this screening after a second round of crosses. If both testis of a pair had the same phenotype, they were scored according to the following criteria: normal bundling, one or more scattered nuclear bundles, no nuclear bundles present, one or more individualization complexes (actin) scattered, no actin cones present, or testis deformed.

### Western Blot

Protein was extracted from 15 WT and α-Spec-GFP fused 3-5 day old flies using NP40 Cell Lysis Buffer (ThermoFisher# FNN0021) + 1mM PMSF + 100X protease inhibitor. Protein was run on BioRad precast gel (BioRad #4568094) Tris/Glycine/SDS in (Bio-Rad# 161-0732), transferred to Immunoblot PVDF membrane (Bio Rad #1620177) in 5X Bolt Transfer buffer (Bio Rad# 10026938). Membranes were incubated at 4°C overnight in blocking buffer (5% BSA + TBST) + antibody (1:2000 dilution); α-Spectrin (DSHB # 3A9 (323 or M10-2)-s), α-Spectrin (Dubreuil Lab), or α-GFP (Cell Signaling Technologies # 2555). Following washes in TBST, membranes were incubated in secondary antibody (Jackson Immuno #115-035-174 or #211-032-171) + 1% BSA (Sigma Aldrich# A7906-500G) (1:5000 antibody dilution) for one hour at room temperature, washed, incubated in Super Signal West Pico Plus Chemiluminescent Substrate (Thermo Scientific #34577), and then imaged. Membranes were stripped after each antibody imaging using Restore Western Blot Stripping Buffer (Thermo Scientific# 21059,).

### Co-Immunoprecipitation Mass Spectrometry

Pairs of testis from 310 3-5 day old flies were dissected for each of 2 replicates of WT and KO flies. Protein was isolated using Pierce IP Lysis Buffer (ThermoFisher# 87788) + Phenylmethanesulfonyl Fluoride (PMSF) + 100X protease inhibitor + SUPERase-In (ThermoFisher# AM2694). All other steps were performed following Sure beads protein G magnetic beads (BioRad# 161-4023) manufacturer’s instructions with monoclonal mouse α-Spectrin antibody (DSHB # 3A9 (323 or M10-2)-s). Protein was then run on BioRad precast gel (BioRad #4568094), excised, and Mass spectrometry was performed by The Taplin Biological Mass Spectrometry Facility at Harvard University. Results from each sample were sorted by number of hits.

Proteins in WT or KO samples that had over 3 hits in control beads samples were excluded. Proteins in WT or KO samples that had fewer than 3 hits in both WT and KO samples were excluded. Ratios of α-Spectrin / protein X were then calculated. All graphs were generated using GraphPad PRISM version 7.03.

### RNA Immunoprecipitation

Testis from 266 WT, KO, or α-Spec-GFP fused fly lines were dissected in ice cold RNase free 1X PBS with tools and surfaces cleaned by RNaseZap RNase Decontamination Solution (Ambion# 9780), fixed in 3% pfa, quenched with 2.5M glycine (Sigma Aldrich# 410225-50G), washed in 1X PBS, flash frozen, and stored at -80. Tissue was lysed in 1000ul Pierce IP lysis buffer (ThermoFisher# 87788) + 10ul PIC + 10ul PMSF + 5ul Superase-in (ThermoFisher# AM2694), incubated on ice for 30 minutes, then centrifuged at 4°C at 16000 x g for 20 minutes. Pull down was performed according to manufacturer’s protocol for Sure Beads protein G magnetic beads (BioRad# 161-4023) and α-Spectrin (DSHB # 3A9 (323 or M10-2)-s), ChromPure Rabbit IgG (Jackson ImmunoResearch Laboratories), or α-GFP (Cell Signaling Technologies # 2555). Following several washes in lysis buffer mix, the beads containing the immunoprecipitated samples were collected and resuspended in 100 ul of: 5ul 1M Tris-CL, pH7.0 (=50 mM) + 1ul 0.5M EDTA (=5 mM) EDTA, + 10ul 0.1 M dithiothreitol DTT (=10 mM) + 5ul 10% SDS (=1%) + 1ul RNase inhibitor SUPERase·in + 78ul H2O RNase-free and crosslinks were reversed by heating at 70^°C^C for 45 minutes. RNA was extracted using Trizol (Life Technologies #15596026) and subjected to DNase treatment (Turbo DNA-free, Ambion# AM1907). Then subjected to cDNA synthesis and PCR.

## Acknowledgements

Special thanks to Dr. Ronald Dubreuil and Dr. Claire Thomas for antibodies, Dr. Melissa Rolls and Dr. Stephen Dinardo for fly lines, the Michael O’Connor Lab for their assistance with their CRISPR protocols, and Dr. Krishanu Ray for spermatogenesis comments. Materials from the *Drosophila* Genomics Resource Center, supported by NIH grant 2P40OD010949.

## Author contributions statement

Conception, Design, Analysis, Interpretation, Manuscript revision: M.B., H.B. Data acquisition, Manuscript writing: M.B. Resources, Funding acquisition: H.B.

## SUPPLEMENTAL INFORMATION

## Supplemental Tables

**Table S1.** Significant SNPs from Lysosome GWAS in relation to lncRNAs -The SNP ID indicates the location of the SNP on the chromosome (2L, 3R, or X). SNP allele identity and p-values were identified and calculated by the Mackay GWAS program^29^. LncRNA names are listed in lncRNA column if the SNP lies within the gene, if not, the following column lists the SNP as “near” the indicated gene followed by the gene’s location on the chromosome according to Flybase version 5.57.

| SNP ID | Minor Allele | Major Allele | Ref Allele | Minor Allele Freq. | Single Mixed Pval | lncRNA | SNP location relative to lncRNA |
| --- | --- | --- | --- | --- | --- | --- | --- |
| 2L_10711046_SNP | T | G | G | 0.05291 | 4.99E-07 |  | near CR44407 ending at 10,710,733 |
| 2L_15399068_SNP | G | C | G | 0.08466 | 6.39E-06 |  | near CR45373 2L:15,398,687..15,399,003 [-] |
| 2L_16948674_SNP | A | G | G | 0.05263 | 1.53E-07 | CR44393 | 2L:16,947,610..16,950,070 [-] |
| 2L_21819676_SNP | A | G | G | 0.05319 | 3.67E-07 | CR44786 | 2L:21,819,490..21,822,313 [+] |
| 2L_21890173_SNP | T | C | C | 0.05263 | 3.32E-06 | CR45362 | 2L:21,889,758..21,891,095 [-] |
| 2L_21890341_SNP | G | A | A | 0.05236 | 2.85E-06 | CR45362 | 2L:21,889,758..21,891,095 [-] |
| 2L_391220_SNP | A | T | A | 0.05263 | 1.37E-06 |  | near CR43718 2L:396,120..396,571 [+] |
| 2L_632462_INS | CC | C | C | 0.05028 | 9.42E-08 |  | near CR45369 2L:628,729..629,052 [+] |
| 2L_632578_SNP | A | G | G | 0.06283 | 9.76E-06 |  | near CR45369 2L:628,729..629,052 [+] |
| 3R_8153667_SNP | C | T | T | 0.05291 | 1.81E-10 | CR45584 | 3R:8,153,069..8,153,725 [+] |
| 3R_8153752_SNP | T | C | C | 0.05729 | 9.74E-06 |  | near CR45584 3R:8,153,069..8,153,725 [+] |
| X_15121154_SNP | T | G | G | 0.09524 | 4.72E-07 |  | near CR44430 X:15,119,546..15,120,289 [+] |
| X_15121172_SNP | C | G | G | 0.09091 | 2.25E-07 |  | near CR44430 X:15,119,546..15,120,289 [+] |
| X_15121280_SNP | T | C | C | 0.08466 | 5.42E-07 |  | near CR44430 X:15,119,546..15,120,289 [+] |
| X_15121286_SNP | G | A | A | 0.07527 | 3.03E-06 |  | near CR44430 X:15,119,546..15,120,289 [+] |
| X_15121307_SNP | G | A | A | 0.08602 | 4.86E-07 |  | near CR44430 X:15,119,546..15,120,289 [+] |
| X_15121308_SNP | G | C | C | 0.08108 | 1.69E-07 |  | near CR44430 X:15,119,546..15,120,289 [+] |
| X_15121329_SNP | G | A | A | 0.07527 | 1.44E-07 |  | near CR44430 X:15,119,546..15,120,289 [+] |

**Table S2.**
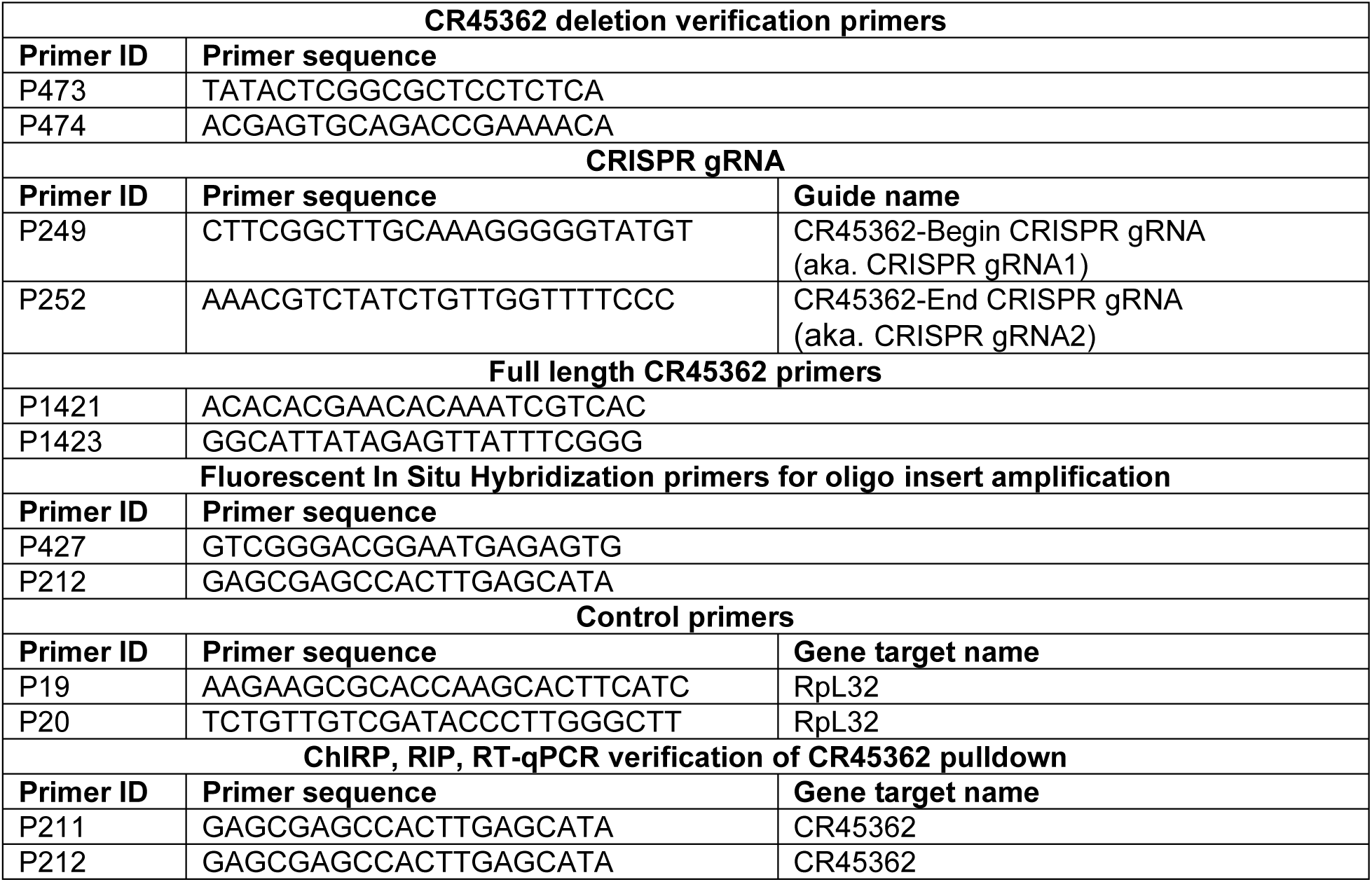
Primer list -Primers with ID# and sequence, sorted by experiment or description.

**Table S3.**
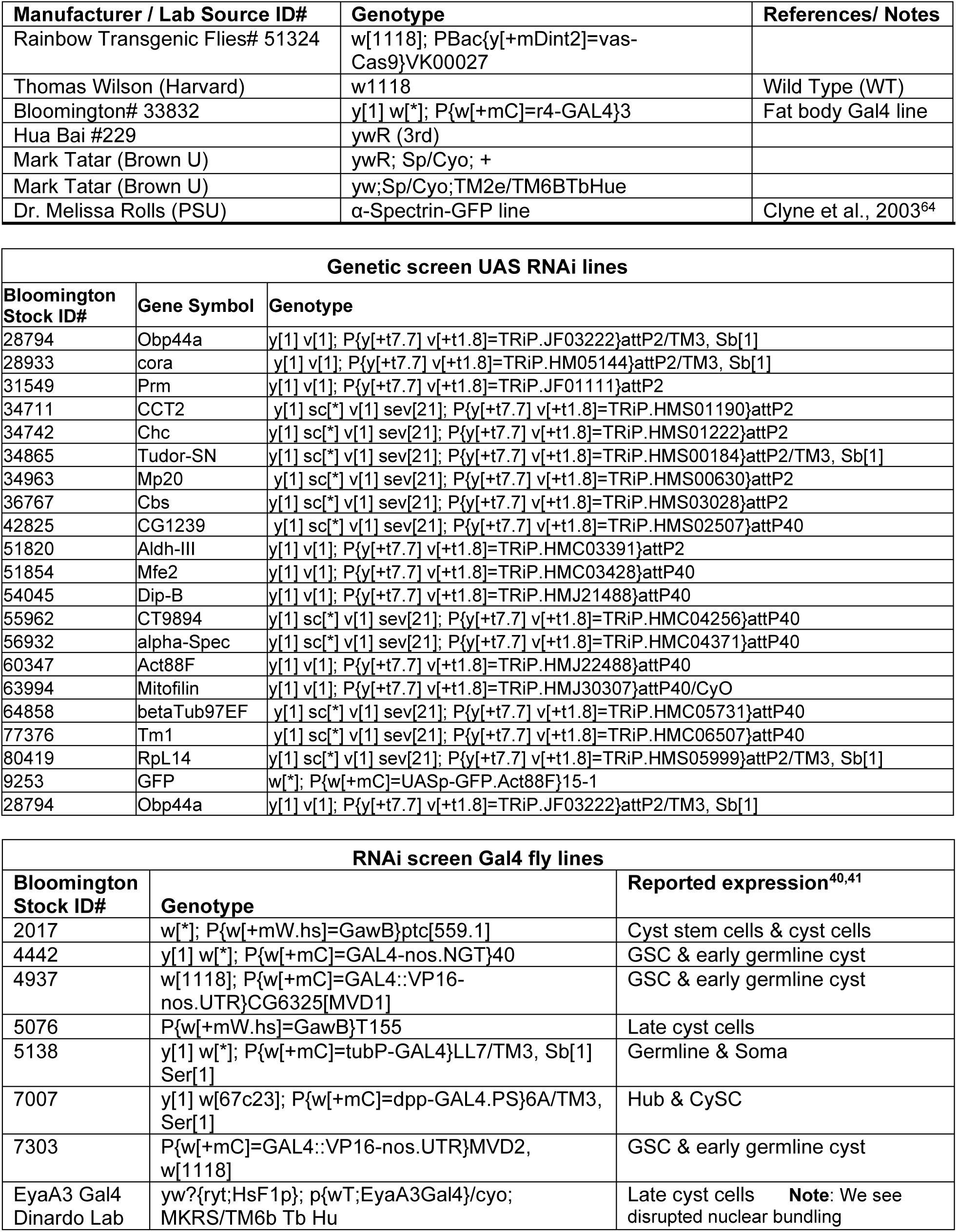
Fly lines -Drosophila melanogaster fly lines used and sources

**Table S4**. Mass spectrometry data

**Supplementary Dataset File:** Excel File: *Mass Spectrometry Data*

Excel Tab1: **ChIRP-MS Run1 Data**

Excel Tab2: **ChIRP-MS Run2 Data**

Excel Tab3: **a-Spec IP-MS Data**

## Supplemental Figures

**Figure S1.**
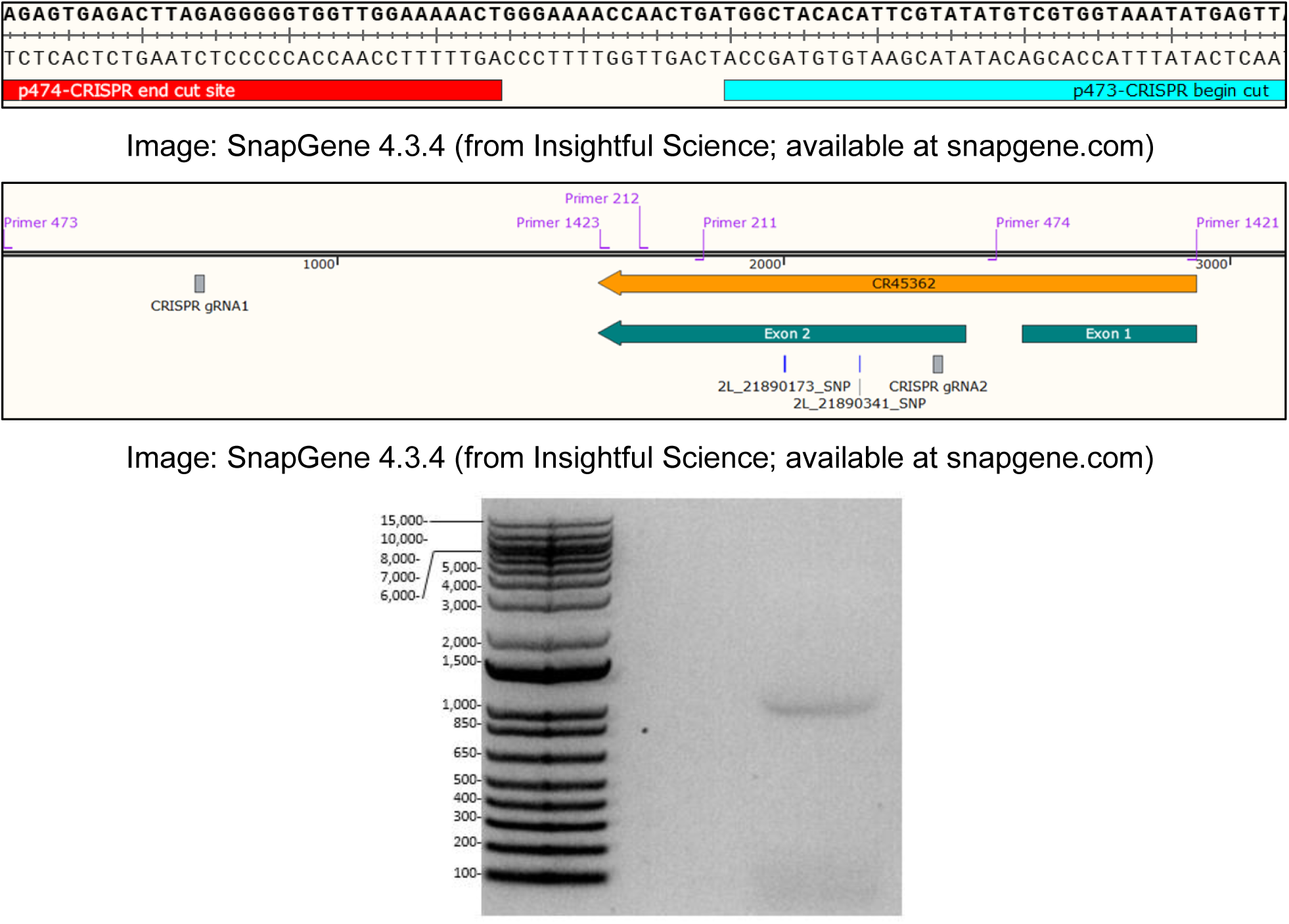
CR45362 CRISPR deletion and full length size verification -Sanger sequencing using primers p474 and p473 (diagrammed in middle box) surrounding the CR45362 deletion loci in the CR45362KO fly mutant (top box) indicates the 2 gRNA target sites have been reduced from 1634 (middle box) to 15 base pairs apart. To check for undescribed isoforms, RNA was extracted from wild type fly lines and converted to cDNA. Using primers 1423 and 1421 (diagrammed in middle box) the cDNA was amplified and run on an agarose gel (bottom box). A single band indicated a single spliced isoform consistent with the 1207bp seen in high throughput sequencing found on Flybase.

**Figure S2.**
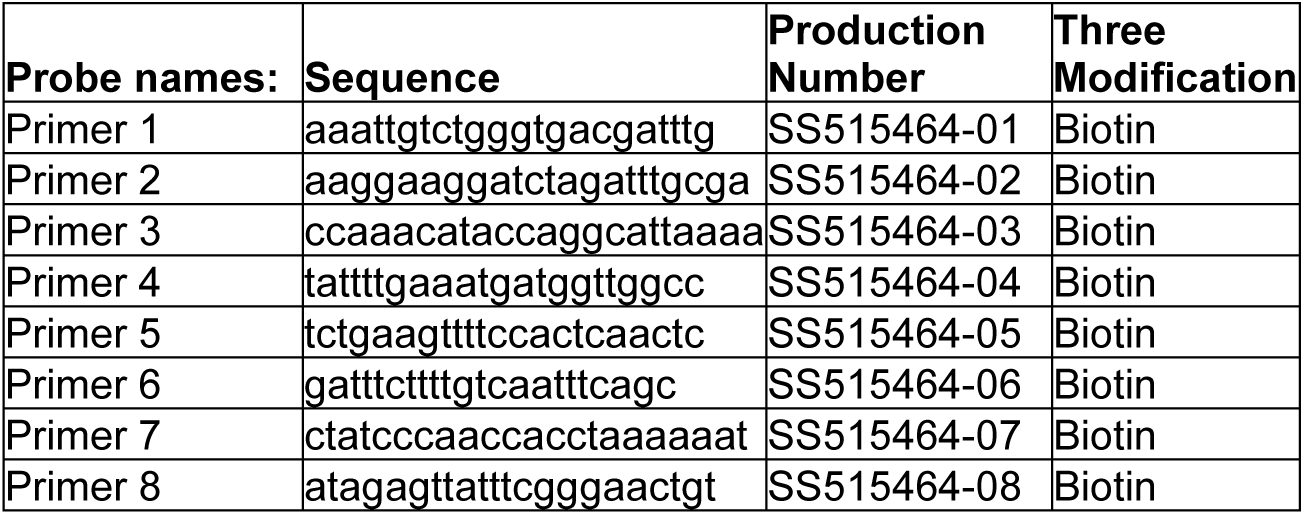
ChIRP probe sequences.-The names and sequences of the 8 probes used in the ChIRP experiment. The production numbers from Bioresearch Technologies are provided for reference and the probe modification was biotin on the 3’ end. Published protocols recommend 1 probe per 200–400 nt of RNA, and avoiding sequences that contain homology to other RNAs.

**Figure S3.**
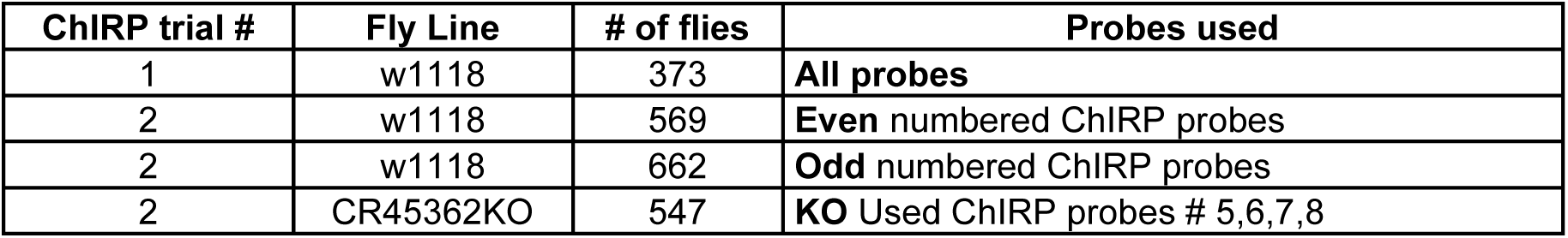
Fly lines, number of flies dissected, and ChIRP probes used for each replicate.

**Figure S4.**
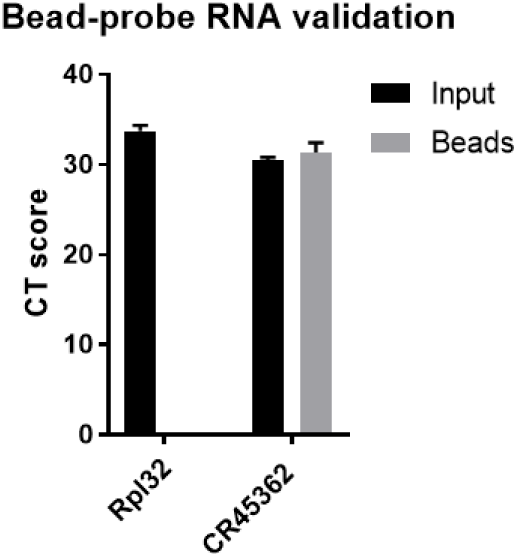
ChIRP RT-qPCR verification of CR45362 pulldown vs control RNA RpL32.-Lysate from WT fly testis was incubated with RNA probes (hybridized to magnetic beads) and then separated. RNA pulled down by probes was compared to RNA from an input sample that was not run on beads. The control ribosomal RNA (Rpl32) is not pulled down by beads, while CR45362 is. Both RNAs are found in Input. Primers 211-212 were used for CR45362 RT-qPCR; values represent 3 technical replicates.

**Figure S5.**
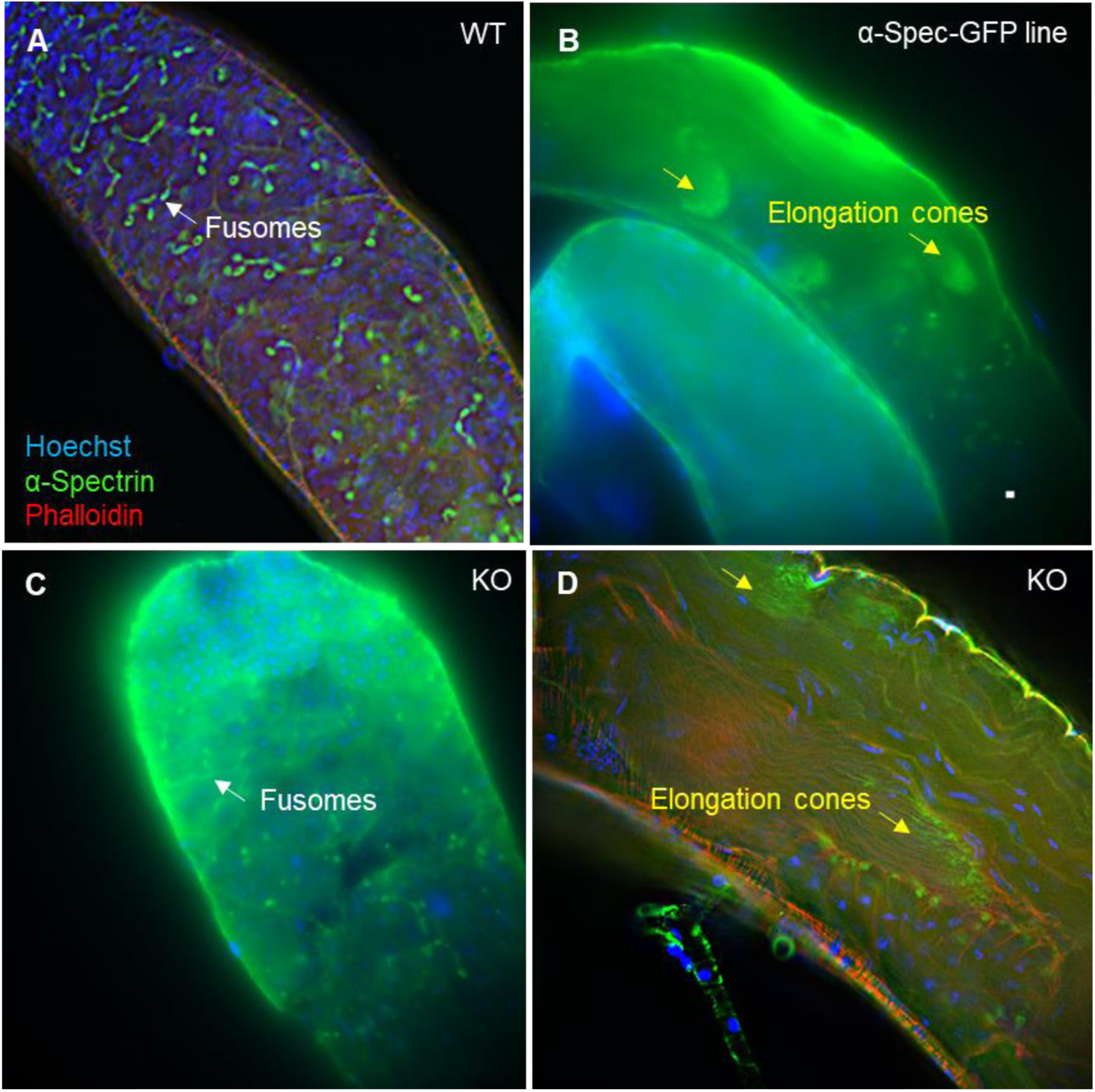
Immunostaining of α-Spectrin structures i.e. fusomes and elongation cones.-The pattern of fusomes (white arrows) in the apical testis of WT flies, indicated by α-Spectrin immunostaining (A), does not appear disrupted in CR45362 mutants (C). Spermatid elongation cones (yellow arrows) in medial testis of α-Spectrin-GFP fused fly line (B), are also present in CR45362 mutants (D).

**Figure S6.**
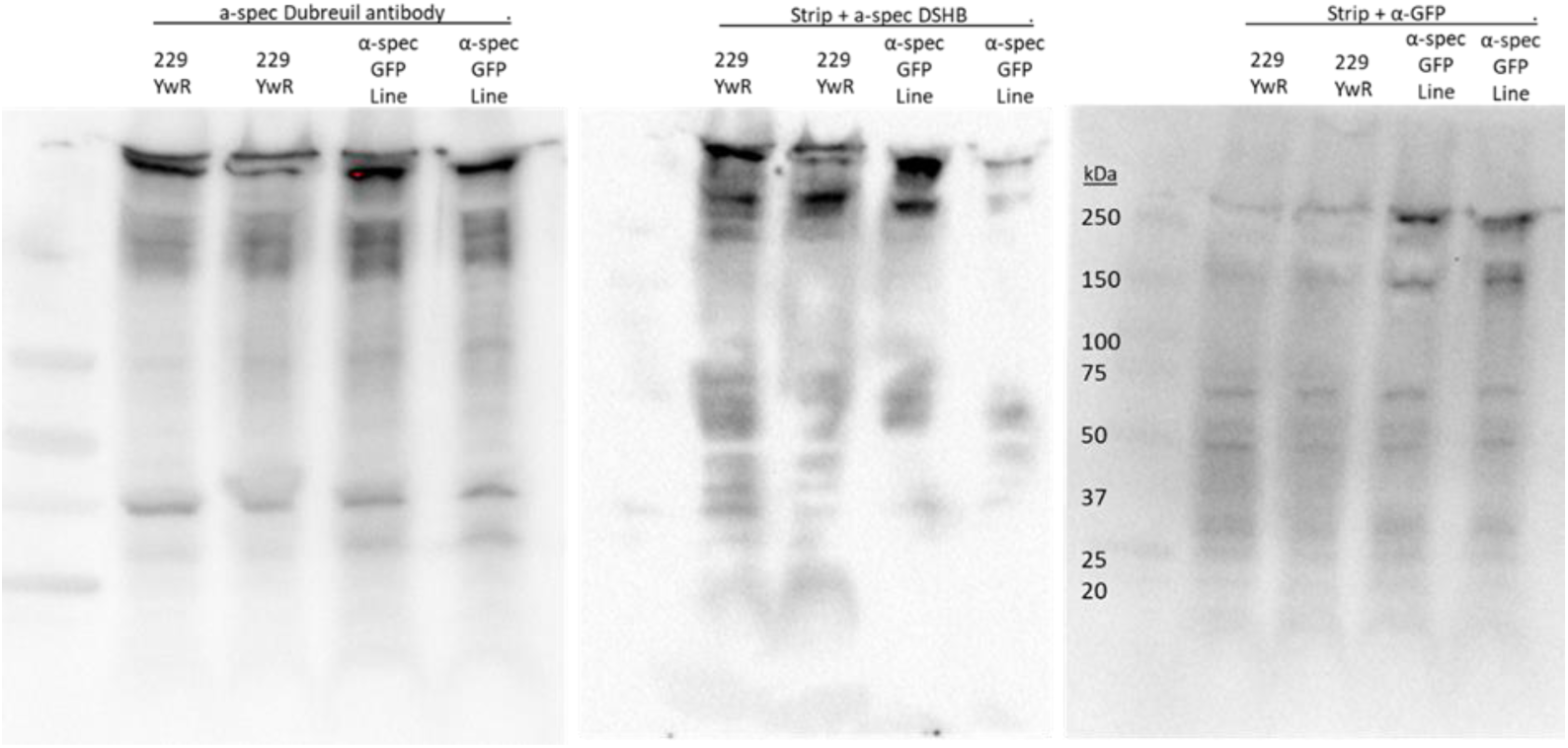
Western blots for α-Spectrin antibodies-An α-Spectrin antibody (a generous gift from Dubreuil lab) was applied to a membrane (left image) against lysate from 2 replicates of WT (YwR) or an α-Spectrin-GFP line. The membrane was stripped and a second α-Spectrin antibody (DSHB) was applied (middle image). The membrane was stripped and an α-GFP antibody applied (right image).

**Figure S7.**
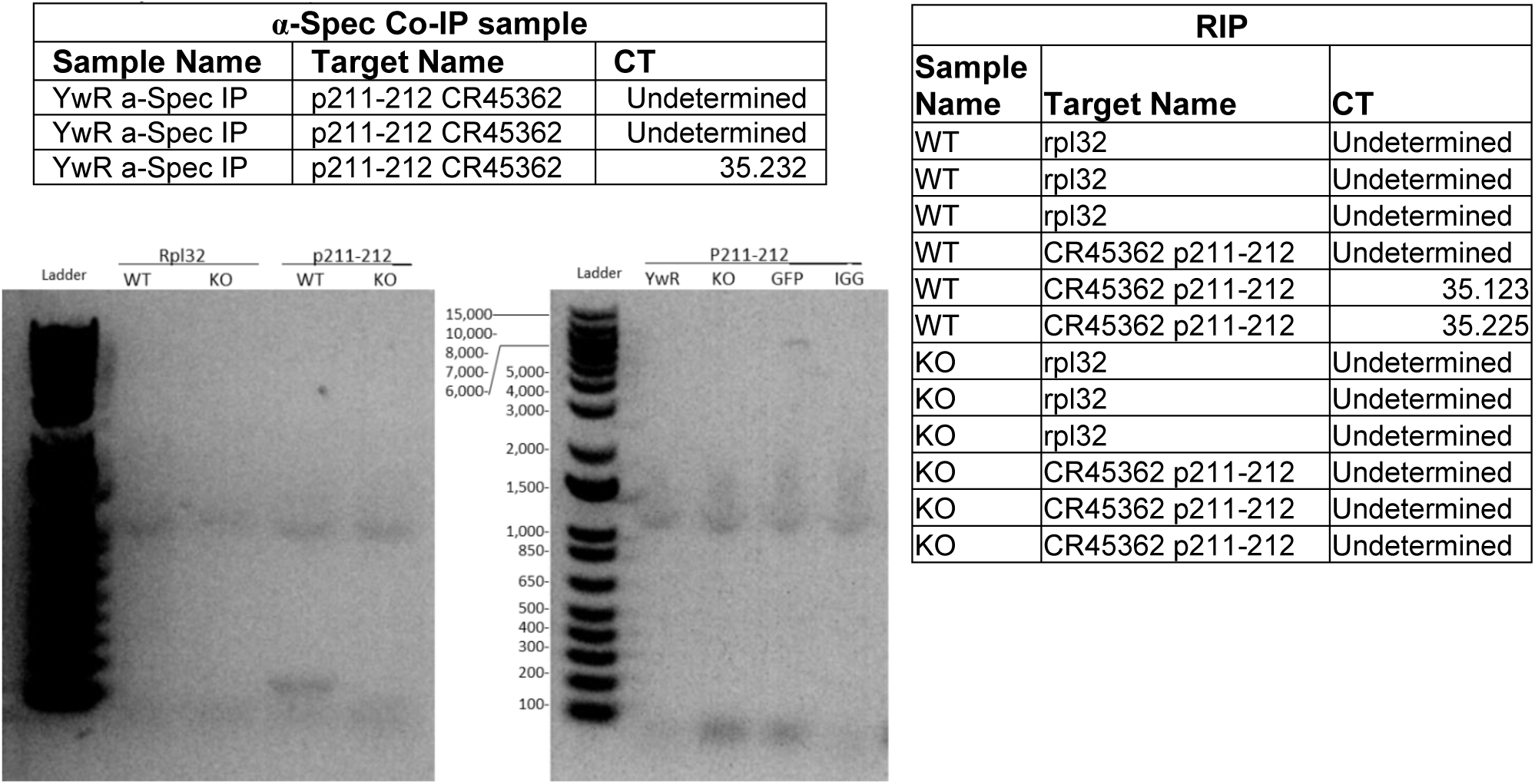
RNA pull-down (Co-IP and RNA Immunoprecipitation)-RNA was purified from the α-Spectrin Co-IP and RT-qPCR was performed (left chart). RT-qPCR was performed on RNA immunoprecipitation pulldown (right chart). CT scores from RIP were confirmed in the left gel indicating the correct product size of 142bp’s using primers p211-p212 was amplified. The right gel indicated that α-GFP and α-IGG did not pull down CR45362 in an α-Spec-GFP fly line.

